# The functional plasticity of the YhdWXYZ ABC transporter enables antibiotic homeostasis and host colonisation in enteric bacteria

**DOI:** 10.64898/2026.05.04.722672

**Authors:** Céline Borde, Géraldine Effantin, Séverine Balmand, Caroline Romestaing, Virginie Gueguen-Chaignon, Agnès Rodrigue

## Abstract

ABC transporters are key determinants of bacterial adaptation, yet their functional plasticity remains poorly understood. Here, we characterize the type I ABC transporter YhdWXYZ and uncover a striking functional divergence linked to the presence of its substrate-binding protein (SBP). In *Escherichia coli* K-12, where *yhdW* is a pseudogene, deletion of *yhdWXYZ* increases susceptibility to mecillinam and lomefloxacin and leads to intracellular accumulation of lomefloxacin, indicating a role in antibiotic homeostasis. In contrast, in *Citrobacter rodentium*, which encodes a complete YhdWXYZ system, deletion of the transporter does not affect antibiotic susceptibility. Biochemical analyses demonstrate that the YhdW SBP of *C. rodentium* binds asparagine with high affinity; however, genetic and physiological assays indicate that YhdWXYZ is not a primary asparagine importer under laboratory conditions, suggesting redundancy with other transport systems. Importantly, *in vivo* infection experiments reveal that YhdWXYZ contributes to early colonization and persistence in the host, as mutants display reduced bacterial loads and altered intestinal pathology in mice. Together, these findings show that loss of the SBP in *E. coli* is associated with a functional shift of YhdWXYZ toward antibiotic homeostasis, whereas in *C. rodentium*, the complete transporter contributes to host adaptation. This work highlights the evolutionary and functional flexibility of ABC transporters in bacterial physiology and pathogenesis.

## INTRODUCTION

ABC (ATP-binding cassette) transporters are present across all domains of life and fulfil diverse transport functions, like the import of essential nutrients and the export of toxic compounds. In bacteria, the number of ABC transporters varies considerably. For example, *Agrobacterium tumefaciens*, a soil bacterium, has more than 200 ABC transporters, while some bacteria have very few (Davidson *et al*, 2008). *Escherichia. coli* K12 has approximately 80 ABC transporters, either experimentally confirmed or presumed (Moussatova *et al*, 2008) (1). The majority are importers of a wide variety of compounds essential to the cell, such as metals, organic ions, amino acids, monosaccharides, oligosaccharides, lipids, or oligopeptides. Bacteria also own ABC transporters that can export toxic compounds. All ABC transporters share a common modular architecture consisting of two highly conserved nucleotide-binding domains (NBDs), and two variable transmembrane domains (TMDs) that form the framework of the translocation pathway. In some importers, a substrate binding protein (SBP) can be found that binds the substrate to be imported with high affinity before transferring it to the TMDs. According to the architecture of the TMDs, ABC transporters are classified into 7 families (Types I-VII) (Thomas *et al*, 2020). All operate solely as importer or exporter, except for type IV transporters which includes importers, exporters, ion channels or regulators (Thomas & Tampe, 2020). Type I are importers associated with an SBP. SBP undergoes a conformational change upon binding of its cognate ligand allowing it to interact with the TMDs and transfer the ligand. Then the translocator applies a second layer of selectivity to exclude non-cognate substrates. In ABC-transporter lacking an SBP, specificity of the transport is secured by the TMDs (de Boer *et al*, 2019).

In this study, we focused on the type I ABC transporter, YhdWXYZ. This transporter consists of an SBP (YhdW), two TMDs (YhdXY) and an NBD (YhdZ). This transporter is conserved in *Escherichia coli* and *Citrobacter spp*., but in *E. coli* K12, a frameshift mutation in the *yhdW* gene results in a truncated SBP protein that may have altered its function. The reason for our interest stems from a previous study in which we demonstrated that the operon encoding the YhdWXYZ transporter is regulated by the ZraSR two-component system (Rome *et al*, 2018). The ZraSR system plays a role in the tolerance of *E. coli* K-12 to some antibiotics, which may be mediated by the YhdWXYZ transporter. To support this hypothesis, a previous study demonstrated the increased susceptibility of the *yhdY, yhdX* and *yhdZ* mutants of *E. coli* K-12 to several antibiotics (Nichols *et al*, 2011). We first tested whether the deletion of *yhdWXYZ* conferred increased susceptibility to certain antibiotics in *E. coli* K-12. We found that its deletion results in susceptibility to mecillinam and lomefloxacin (a fluoroquinolone), as well as higher intracellular accumulation of lomefloxacin. This suggests that this transporter plays a role in antibiotic tolerance in *E. coli* K12. To study an orthologous system with a functional YhdW, we selected the murine pathogen bacterium *Citrobacter rodentium*. In this bacterium, we show that YhdWXYZ is not associated with antibiotics susceptibility, and that purified YhdW binds asparagine with high affinity. Finally, we demonstrate that YhdWXYZ from *C. rodentium* plays an important role in host-pathogen interaction.

## MATERIALS AND METHODS

### Bacterial strains and growth conditions

The strains used in this work are derivatives of *E. coli* K-12 or *C. rodentium* DBS100. Bacterial cells were grown at 37°C in LB medium or M63 minimal medium. Antibiotics and amino acids were purchased from Merck. Stains and plasmids used in this study are listed in Table S1 and S2.

### Deletion mutant construction

Yhd gene knockouts were created using the lambda red recombination technique (Datsenko & Wanner, 2000) in *E. coli* or an adapted protocol (Crepin *et al*, 2015) in *C. rodentium*. Deletion PCR products containing upstream and downstream homologous regions of 50 bp for *E. coli* or 300 bp for *C. rodentium* were generated using pKD3. *E. coli* or *C. rodentium* harboring pKD46 or pKOBEG respectively were used to perform recombination and resistance cassettes were removed using pCP20. Deletions were verified by PCR and sequencing.

### Core-genome phylogeny and Yhd proteins conservation

Genome accession numbers of *Escherichia spp., Shigella spp. Citrobacter spp.* and *Salmonella spp.* included in this study are provided in Table S3. *Klebsiella* and *Enterobacter* were used as outgroups. Genomes were annotated with Prokka v1.14.6 using default parameters to generate standardized GFF3 annotation files for comparative genomic analyses. Annotated genomes were analyzed using the pipeline Roary v3.13.0 to identify orthologous gene clusters and define core and accessory gene sets. A BLASTP percentage identity threshold of 90% was applied for gene clustering, and core genes were defined as genes present in 99% of the analyzed isolates. The Roary-generated core gene alignment was subsequently used for phylogenetic reconstruction. A maximum-likelihood phylogenetic tree was inferred from the concatenated core-genome nucleotide alignment using IQ-TREE with automatic substitution model selection (ModelFinder). Branch support was assessed using 1,000 ultrafast bootstrap replicates. The resulting tree was visualized and annotated using Interactive Tree Of Life (iTOL).

To investigate the distribution and structural conservation of the *yhdWXYZ* genes, each gene was used as queries in BLASTn searches against all analyzed genomes using Geneious. Gene presence or absence was determined based on sequence similarity thresholds including percentage identity, alignment coverage, and E-value. In addition to gene detection, BLASTn results were manually inspected to identify sequence alterations affecting the locus, including frameshift mutations, insertion sequence (IS) element insertions, and other gene-disrupting events. The resulting gene conservation and structural variation data were manually mapped onto the core-genome phylogenetic tree to visualize the evolutionary distribution of the system and to compare gene integrity patterns across *Escherichia, Shigella, Citrobacter* and *Salmonella*.

### Antibiotics susceptibility assays

#### Challenge test

Each strain was inoculated into fresh LB medium and grown to an OD600 of 0.1 at 37°C. Cultures were challenged with the indicated antibiotics at the concentrations listed in the Figure 1 legend. Cells were incubated in microtubes at 37°C with shaking at 550 rpm for 1 h and then plated on LB agar plates overnight at 37°C, as described previously (Rome et al., 2018). Survival percentage was calculated as follows: survival (%) = CFU (colony-forming units) of surviving cells / CFU of non-challenged cells. The relative survival rate of the mutant (Δ*yhdW*-Z) was calculated as the ratio of survival percentages (WT/mutant).

**Figure 1.**
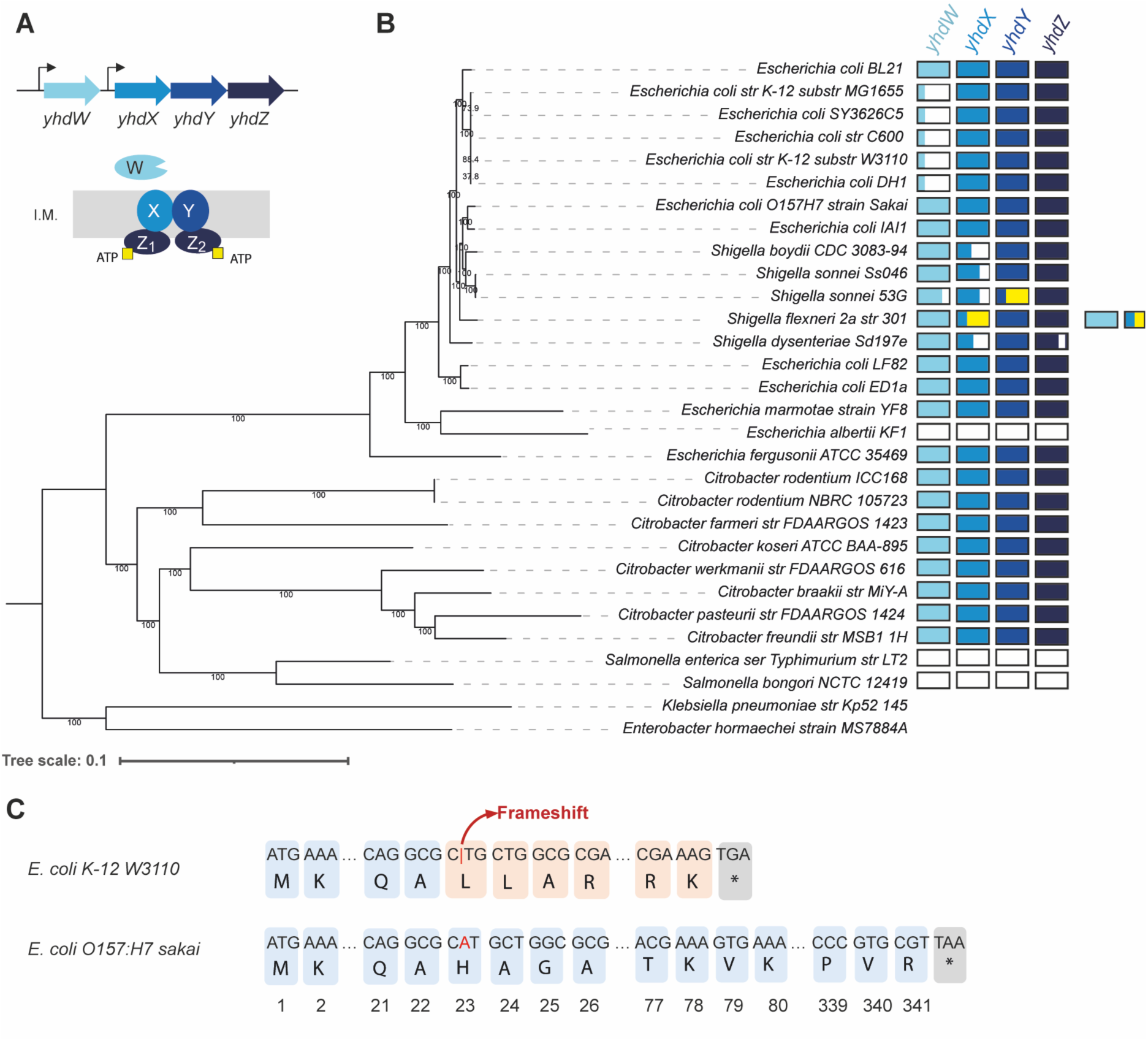
: Distribution and genetic variations of *yhdWXYZ* genes among *Enterobacteriaceae*. **(A)** Predicted organization of the YhdWXYZ ABC transporter. **(B)** A maximum likelihood phylogenetic tree derived from the nucleotide alignment of the concatenated central genome using IQ-TREE. The presence or absence, as well as the structural status, of each gene belonging to the Yhd system has been plotted on the tree. Blue boxes indicate intact genes present in the corresponding genome, whilst empty boxes indicate the absence of a gene. Blue and white boxes indicate pseudogenised genes (frame-shift resulting in truncated coding sequences) and blue and yellow boxes indicate genes interrupted by insertion sequence (IS) elements. Partial duplication of the locus is further indicated in the annotation track for *S. flexneri* 2a str 301. This analysis highlights the heterogeneous conservation and lineage-specific modifications of the system in *Escherichia, Shigella, Citrobacter* and *Salmonella*. **(C)** In *E. coli K-12*, the deletion of a single A base in the coding region causes a frameshift starting at amino acid 23 and the appearance of a premature stop codon.

#### Growth assays in the presence of subinhibitory antibiotic concentrations

The pvlt33-yhdWXYZ plasmid was constructed by inserting the PCR-amplified *yhdWXYZ* operon from *E. coli* using the Primestar Max (Takara) into the pvlt33 vector digested with *Sac*I/*Xba*I using the TLTC method (Yu *et al*, 2023). Bacteria were grown at 37°C in 96-well plates in LB supplemented with1 mM IPTG, or in LB supplemented with IPTG 1 mM and either lomefloxacin (0.075 µg/mL) or tetracycline (0.2 µg / mL) for *E. coli* K-12; or in LB supplemented with lomefloxacin (0.25 µg/mL) or tetracycline (0.2 µg / mL) for *C. rodentium*. The O.D._600nm_ was measured every 20 min for 24 hours in the Infinite® 200 PRO plate reader (Tecan).

#### Promoter activity assays

To measure the activity of the *yhdW* or *yhdX* promoters in *E. coli* and *C. rodentium*, transcriptional fusions were constructed by cloning the PCR-amplified promoters regions upstream of the *luxCDABE* operon in the pLux plasmid (Cayron *et al*, 2017). Assays were performed in 96-well plates containing 200 µL of LB medium inoculated with bacteria harvested in exponential phase (OD600 = 0.6). Plates were incubated at 37°C with shaking (280 rpm) for 24 to 48 hours in the Infinite® 200 PRO plate reader (Tecan). OD600 and luminescence were measured every 20 min following a 1 min shaking period.

#### Intracellular lomefloxacin quantification

The cell preparation protocol was adapted from the publication by Vergalli et al. (Vergalli *et al*, 2018). Bacteria were grown in rich medium LB to an OD600 of 0.6 and collected by centrifugation. Cells were resuspended in NaPi-Mg buffer (NaH_2_PO_4_ 44.2%, 55.8 NaHPO_4_, and MgCl_2_ 0.25%) to a final OD600 of 6. Cells were then incubated at 37°C with or without 1μg/mL lomefloxacin. Aliquot were collected after 10, 20, 30 or 45 min. Cell numbers were determined by dilution and plating, and cells were centrifuged on a 1M sucrose cushion. The supernatant was removed and cells were lysed in a 200μL lysis buffer (50% acetonitrile and 2.5% trifluoroacetic acid). Cell debris was removed by centrifugation and supernatants were analyzed.

The HPLC system consisted of a Waters 2795 Alliance HT separation module, and detection was performed using a Waters 2996 photodiode array detector coupled to an SFM25 fluorometer (Kontron). Samples were separated on a Gemini 5-micron C18 column (250 X 4.6 mm, Phenomenex) at 40°C with a flow rate of 1mL/min. Elution was performed using a linear gradient: 0–5 min, 100% A; 5–15 min, 55% A and 45% D; 15–25 min, 0% A and 100% D; 25–30 min, return to 100% A, followed by 5 min re-equilibration. Solvent A contained 0.005% trifluoroacetic acid, and solvent D consisted of 100% methanol.

The emitted fluorescence was measured with excitation at 280 nm and emission at 440 nm. Lomefloxacin was quantified by measuring peak areas using chromatographic data processing software (Mass Lynx, Waters). A standard curve was used to correlate fluorescence intensity with lomefloxacin concentration. The intracellular amount of lomefloxacin was calculated by dividing the total amount by the number of cells.

#### Overproduction and purification of the *C. rodentium* YhdW protein

To produce and purify YhdW from *C. rodentium*, the *yhdW* open reading frame was cloned upstream of an 8×His tag in the IPTG-inducible pET30 vector. *E. coli* BL21 cells carrying the pET30-YhdW-HT plasmid were grown in LB at 30°C to an OD600 of approximately 0.3 and induced with 0.1 mM IPTG for 4 h. Periplasmic proteins were extracted by osmotic shock. Cells were resuspended in 50 mM Tris-HCl (pH 8) containing 1 mM EDTA, followed by addition of 40% sucrose. After incubation on ice and centrifugation, cells were resuspended in water containing 1.25 mM MgSO_4_ and incubated again on ice. The periplasmic fraction was collected and incubated with Ni-NTA beads (Qiagen) in the presence of 10 mM imidazole, 1 mM DTT, and protease inhibitors (Roche).

The mixture was loaded onto a column, washed with buffer A (10 mM Tris-HCl pH 8, 100 mM NaCl, 5% glycerol, 2 mM β-mercaptoethanol) supplemented with 10 mM and then 30 mM imidazole, and eluted with buffer A containing 150 mM imidazole. Fractions were analyzed by SDS-PAGE, and the purified protein showed no detectable contaminants. Imidazole was removed by dialysis using PD-10 desalting columns (Cytiva).

#### Fluorescence Thermal Shift Assay

FTSA was performed using a Mosquito nanopipetting robot (SPT Labtech) to mix purified YhdW, SYPRO Orange dye (Molecular Probes Thermofisher), and amino acids where indicated, in a final volume of 25 µL buffer A in 96 wells plate (MicroAmp Fast Optical Reaction Plate (Thermofisher)). A Real-Time PCR system StepOne Plus (Applied Biosystems) was then used to create a temperature gradient from 25°C to 95°C at 1°C per minute, and to measure the fluorescence intensity at 530 nm after excitation at 485 nm. Initial screening was performed with 1.5 µM protein in the presence or absence of 20 amino acids at concentrations of 0.2 mM or 1 mM. A second screening used 2.5 µM protein with amino acids at concentrations ranging from 0.25 to 1 mM. Melting temperatures (Tm) were determined from the first derivative of fluorescence data. A ΔTm greater than 2°C was considered indicative of binding.

#### Isothermal titration calorimetry (ITC)

ITC measurements were performed using a MicroCal ITC200 microcalorimeter at 25°C in 10 mM Tris-HCl (pH 8), 5% glycerol, 1 mM β-mercaptoethanol, 75 µM imidazole, and 0.5 µM DTT. Approximately 200 µL of protein (200 µM) was placed in the cell and titrated with a 10× concentrated amino acid solution. After an initial 2 µL injection, 14 injections of 4 µL were performed at 2 min intervals. Data were corrected for heat of dilution and analyzed using a single-site binding model in ORIGIN software. The binding enthalpy change (ΔH), association constant (Ka), and binding stoichiometry (n) were permitted to float during the least-squares minimization process and taken as the best-fit values.

#### Growth competition

Cells from LB precultures were washed in M63 medium and adjusted to identical OD600 values. A 1:1 mixture of *ΔasnAΔasnB* and *ΔasnAΔasnBΔyhdWXYZ*-KanR strains was inoculated at an OD600 of 0.05 in M63 medium supplemented with 0.2% glucose and either 0.1 mM, 0.5 mM or 1 mM asparagine. Cultures were incubated at 37°C with shaking. At 3, 6, and 24 h, samples were plated on LB and LB supplemented with kanamycin. The proportion of triple mutants was calculated (N = 4).

#### Animal care and housing

Experiments were approved by the Lyon ethics committees and the Ministry of Research and Higher Education (Ministère de la Recherche et de l’Enseignement Supérieur), and conducted in accordance with animal care guidelines (APAFIS#9997-2017052213468524v2).

Six-week-old female C57BL/6 mice were supplied by Charles Rivers, France. All animals were housed in an experimental animal facility (“ACSED”, University of Lyon 1, France), under a 12 h light/dark cycle, in a temperature-controlled room and with food and water provided *ad libitum*. Animals were randomly assigned into cages upon reception. After acclimatation, cages where randomly assigned to treatment groups (without (control) or with bacterial suspensions of *C. rodentium* WT, Δ*yhdW*, or Δ*yhdWXYZ* strains (n = 9 per group)).

#### Mouse infection

Infection experiments were performed as described previously (Crepin *et al*., 2015). Bacterial cultures were grown overnight, centrifuged, and resuspended in PBS. Mice were orally inoculated with 200 µL of bacterial suspension or PBS for controls. Bacterial loads in feces were determined daily by plating serial dilutions on MacConkey sorbitol agar.

At days 7 and 12-14 post-infection, mice were euthanized by cervical dislocation in accordance with scientific procedures for animals (Act of 1986) and local ethical guidelines. Colon samples were collected, fixed in 4% paraformaldehyde, and dehydrated using graded concentrations of ethanol prior to impregnation and embedding in Paraplast®. Microtome sections were prepared and processed for histological analysis (HE (hematoxylin and eosin) or PAS (Periodic Acid-Schiff) staining. Crypt hyperplasia was quantified by measuring the length of 10 well-oriented crypts per sample using QuPath software.

## RESULTS

### The YhdWXYZ transporters shows variable conservation and frequent genomic alterations across *Enterobacteriaceae*

YhdWXYZ belongs to the type I ABC transporter family, in which YhdW is the predicted SBP, YhdX and YhdY are predicted permeases, and YhdZ is the ATPase (Figure 1A). YhdZ belongs to the K09972 KEGG orthology group, corresponding to general L-amino acid transport ATP-binding proteins. Homologs are widely distributed across bacterial phyla, including Proteobacteria, Firmicutes, Actinobacteria, and others. To assess conservation of the YhdWXYZ transporter, the distribution and integrity of the *yhd* genes were displayed on a phylogenetic tree inferred from the core genome alignment of Enterobacteriaceae (Figure 1B). The *yhdWXYZ* system was consistently detected in *Escherichia coli, Shigella spp*., and *Citrobacter spp.,* whereas it was absent from *Escherichia albertii* and *Salmonella spp.*. This absence may reflect gene loss events.

In *Citrobacter spp.* and *E. coli*, the system was found to be generally well conserved, with high sequence identity and a genetic organization that is preserved in most strains. However, a frameshift mutation corresponding to a deletion at nucleotide 68 was identified in the *yhdW* gene of the *E. coli* K-12 strains, resulting in a predicted 78-amino-acid truncated protein (Figure 1C). This protein will no longer encode a functional SBP in *E. coli K-12* related strains and doesn’t display any homology with other ORFs. Similarly, *Shigella* genomes exhibited a higher level of structural variation within the system. Multiple insertion sequence (IS) elements were detected within the four genes, along with frameshift mutations and rearrangements. Furthermore, a genetic duplication event was observed at one locus, suggesting ongoing genomic remodeling.

Overall, these observations indicate that while the system is largely conserved within *Citrobacter* and *Escherichia*, it shows signs of degeneration and structural rearrangement in *Shigella spp* and *E. coli* K12, which could indicate a diversification of the transporter’s function.

### The *yhdWXYZ* mutant shows increased sensitivity to specific antibiotics in *E. coli* K-12

We previously demonstrated that deletion of the ZraSR two-component system (TCS) increases susceptibility to various antibiotics in *E. coli*. The ZraSR TCS regulates the expression of several genes, including the *yhdWXYZ* operon encoding an ABC transporter (Rome *et al*., 2018). To determine whether YhdWXYZ contributes to the antibiotic response mediated by ZraSR, the Δ*yhdWXYZ* mutant was exposed for one hour to a range of antibiotics from different families, and survival was assessed.

Exposure to tetracycline, puromycin, and ampicillin resulted in similar survival rates between the wild-type (WT) strain and the Δ*yhdWXYZ* mutant. In contrast, treatment with lomefloxacin or mecillinam led to a significant reduction in mutant viability compared to the WT strain (Figure 2A). To further evaluate these effects, growth of the WT strain, the Δ*yhdWXYZ* mutant, and the complemented strain was monitored in the presence of subinhibitory concentrations of lomefloxacin or tetracycline. Consistent with the survival assays, tetracycline had a comparable impact on all strains (Figure 2B). In contrast, lomefloxacin had a significantly stronger inhibitory effect on the mutant strain. This growth defect was partially complemented by expression of the *yhdWXYZ* operon from a plasmid (Figure 2B). Complete growth curves are provided in Supplementary Figure S1.

**Figure 2:**
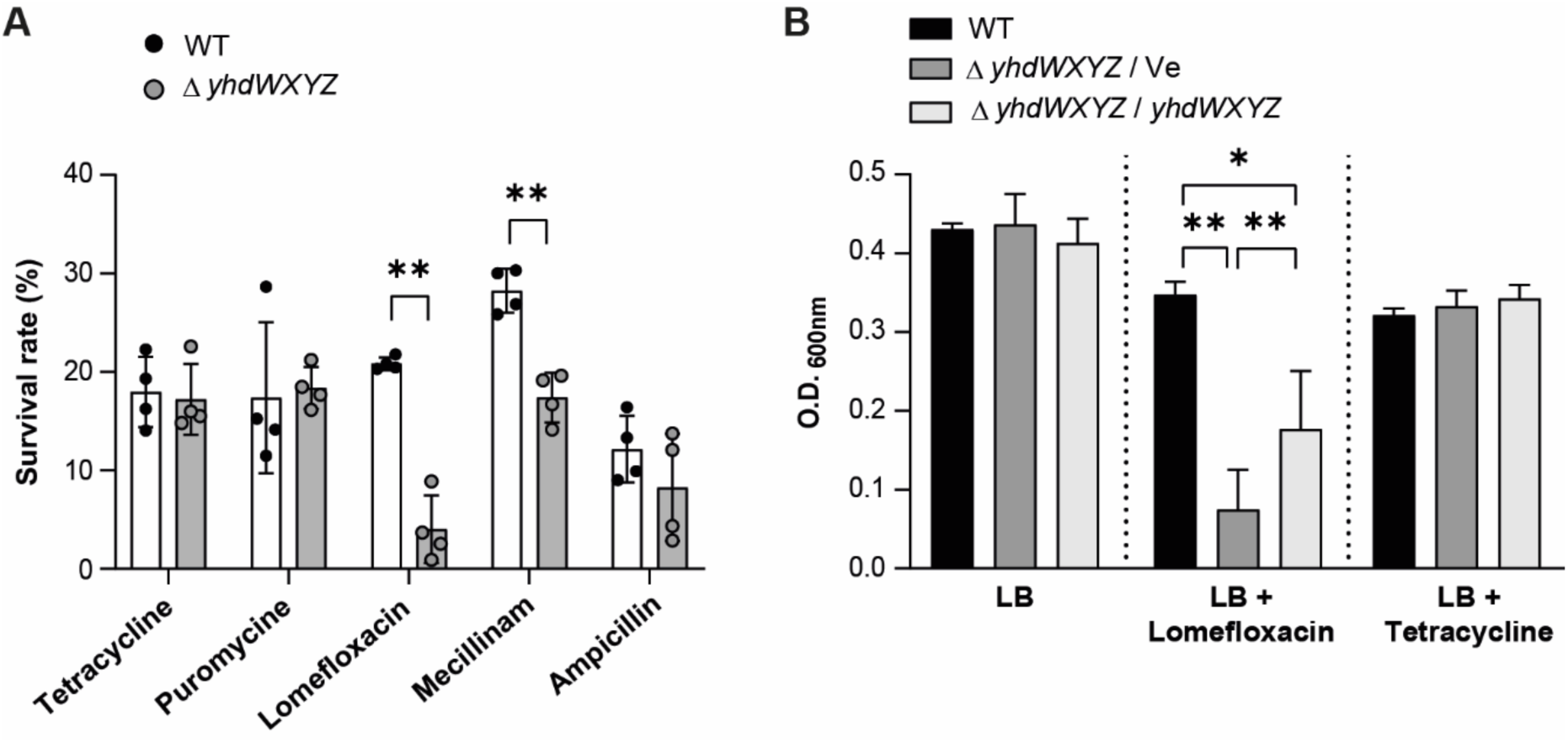
*yhdWXYZ* deletion confers susceptibility to some antibiotics in *E. coli* K-12. **(A)** Screening test. Bacteria were plated after being in contact with subinhibitory concentrations of : Tetracycline (35 µg/mL), Puromycine (35 µg/mL), Lomefloxacin (0.2 µg/mL), Mecillinam (5 µg/mL) or Ampicillin (10 µg/mL). The mean and SD of 4 replicates are shown, pVal < 0.01, 2 way ANOVA. **(B)** Growth of the WT strain, the Δ*yhdWXYZ* mutant / vector and the Δ*yhdWXYZ* / *yhdWXYZ* strain. The bacteria were grown), at 37°C in LB + IPTG 1mM, or in LB + IPTG 1mM + Lomefloxacin (0.075 µg/mL) or Tetracycline (0.2 µg / mL). The O.D.600nm was measured for 24 hours. The values after 15h of growth are shown. The mean and SD of 7 replicates are shown, Kruskal Wallis test.

### Deletion of *yhdWXYZ* leads to increased intracellular accumulation of lomefloxacin in *E. coli* K-12

YhdWXYZ belongs to the type I ABC transporter family, in which *yhdW* no longer encodes a functional SBP (Figure 1C). As type I ABC transporters can also function as exporters, we investigated whether YhdWXYZ contributes to lomefloxacin export. Lomefloxacin was added to non-growing bacterial cells, which were sampled at regular intervals. For each sample, colony-forming units (CFUs) were determined, and intracellular lomefloxacin levels were quantified by HPLC.

In WT cells, intracellular lomefloxacin levels remained relatively constant over time. In contrast, the Δ*yhdWXYZ* mutant showed a progressive increase in intracellular lomefloxacin (Figure 3A), reaching approximately twice the level observed in the WT. Increased variability at later time points likely reflects higher cell mortality in the mutant population. These results indicate that YhdWXYZ contributes to the homeostatic control of intracellular lomefloxacin levels.

**Figure 3:**
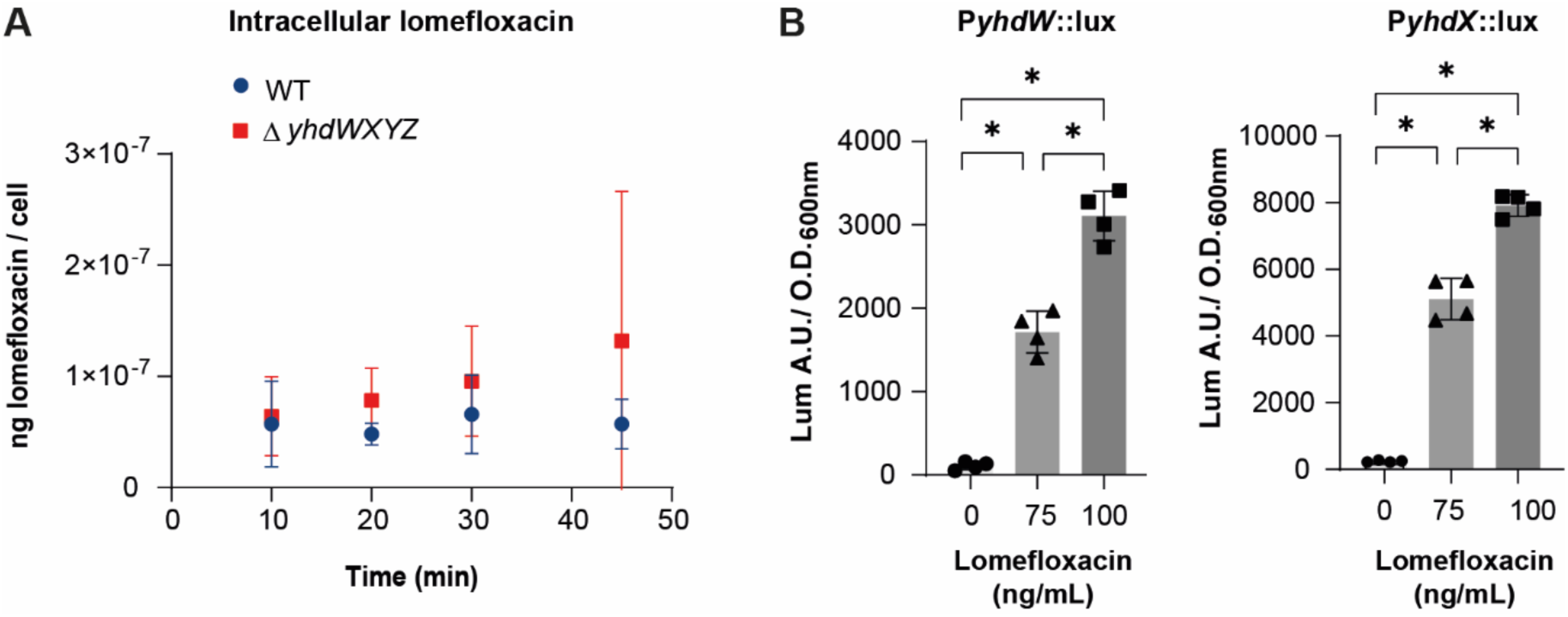
Deletion of *yhdWXYZ* promotes lomefloxacin accumulation in *E. coli* K-12. (**A)** Lomefloxacin was added to non-growing bacteria. Samples were collected every 10 minutes, and the intracellular concentration of lomefloxacin was measured using HPLC. The mean and standard deviation for four repetitions are shown. **(B)** Expression of the *yhdW* (left) and *yhdX* (right) promoters in presence of increasing concentrations of Lomefloxacin in LB medium. Plasmid borne *lux* fusions were assayed in the WT strain, and luminescence specific activity recorded. The mean and SD of 4 replicates are shown, pVal < 0.05, Mann-Whitney test.

To further investigate this hypothesis, expression of the *yhdWXYZ* operon was analyzed. Two promoters are predicted: one upstream of *yhdW* and another upstream of *yhdX*. Transcriptional fusions of these promoter regions to the *luxCDABE* reporter were constructed, and luminescence was measured in the presence of increasing concentrations of lomefloxacin. Both promoters were induced in a dose-dependent manner by lomefloxacin (Figure 3B), supporting a role for YhdWXYZ in antibiotic homeostasis in *E. coli* K-12.

### YhdWXYZ is not involved in antibiotic susceptibility in *C. rodentium*

To investigate the role of a complete YhdWXYZ transporter, we analyzed its function in *C. rodentium*, which encodes a full system including a functional SBP. Sequence comparison showed high conservation with *E. coli* proteins (YhdX: 85% identity; YhdY: 89%; YhdZ: 92%). Moreover, we wanted to find a suitable model to assess its role *in vivo*. To study closely related non-human pathogenic bacteria, we selected *C. rodentium*, a rodent pathogen that is harmless to humans. *C. rodentium* is a model bacterium for studying *E. coli* infections, as the virulence mechanisms involved in host interactions rely on conserved molecular patterns (Collins *et al*, 2014) (see below).

Antibiotic susceptibility assays were performed as described for *E. coli*. In contrast to the results obtained in *E. coli* K-12, the Δ*yhdWXYZ* mutant of *C. rodentium* did not display any significant growth defect compared to the WT strain in the presence of subinhibitory concentrations of lomefloxacin or tetracycline (Figure 4). These results indicate that the presence of the SBP alters the functional role of the transporter. The complete growth curves are shown in supplementary data (Figure S2).

**Figure 4:**
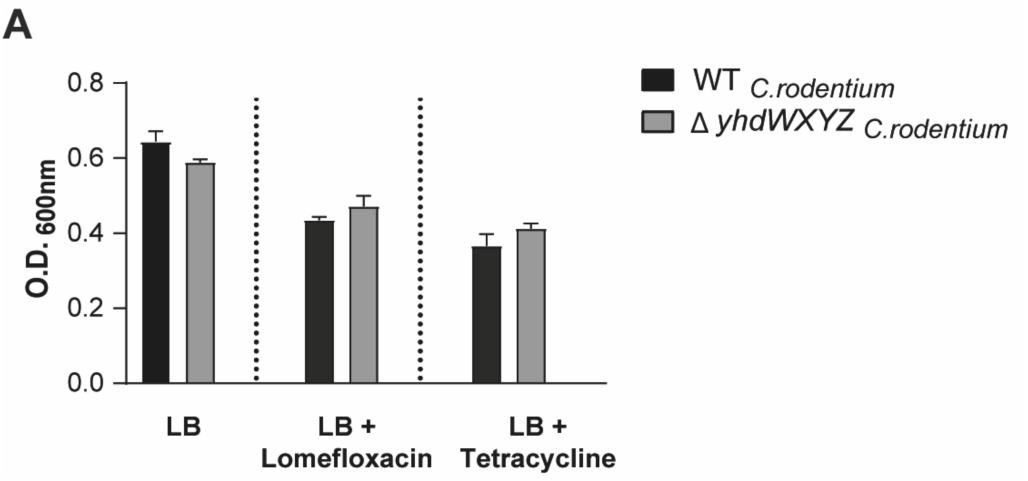
The full YhdWXYZ transporter of *C. rodentium* is not involved in antibiotics sensitivity. (A) CMI of *C. rodentium* RCL2 WT strain, the *ΔyhdWXYZ isogenic* mutant / vector and the *ΔyhdWXYZ mutant* / complemented by *yhdWXYZ* of *E. coli* K-12. The bacteria were grown in LB + IPTG 1mM, or in LB +1mM IPTG + subinhibitory concentrations of : Lomefloxacin (0.25 µg/mL) or Tetracycline (0.2 µg / mL). The O.D._600nm_ was measured for 24 hours at 37°C. The values after 15h of growth are shown. The mean and SD of 5 replicates are shown.

### The YhdW SBP from *C. rodentium* is an asparagine-binding protein

The Alphafold server (powered by Alphafold 3, (Abramson *et al*, 2024)) was used to model the folding of each protein of the YhdWXYZ ABC-transporter. Then, the predicted 3D structures were compared with the RCSB Protein Data Bank (PDB) using the DALI server (Holm *et al*, 2023). YhdW_Crod_ best structural neighbor is BztA, a periplasmic binding protein of an ABC-transporter from *Brucella ovis* (PDB: 4z9n, unpublished structure). This protein has not been characterized in this bacterium, however, its ortholog in *Rhodobacter capsulatus* was shown to transport glutamate, glutamine, aspartate and asparagine (Zheng & Haselkorn, 1996). YhdW_Crod_ was cloned with an 8xHis C-terminal tag, overexpressed and purified from the periplasmic fraction (Figure S3).

To determine ligand specificity, purified YhdW was subjected to fluorescence thermal shift assays (FTSA) in the presence of amino acids. Among the 20 amino acids tested, asparagine produced the largest increase in melting temperature (ΔTm), indicating strong binding (Figure 5A). Secondary screening confirmed that asparagine induces a significant concentration-dependent increase in Tm, whereas other amino acids showed minimal or no effect (Figure 5B).

**Figure 5.**
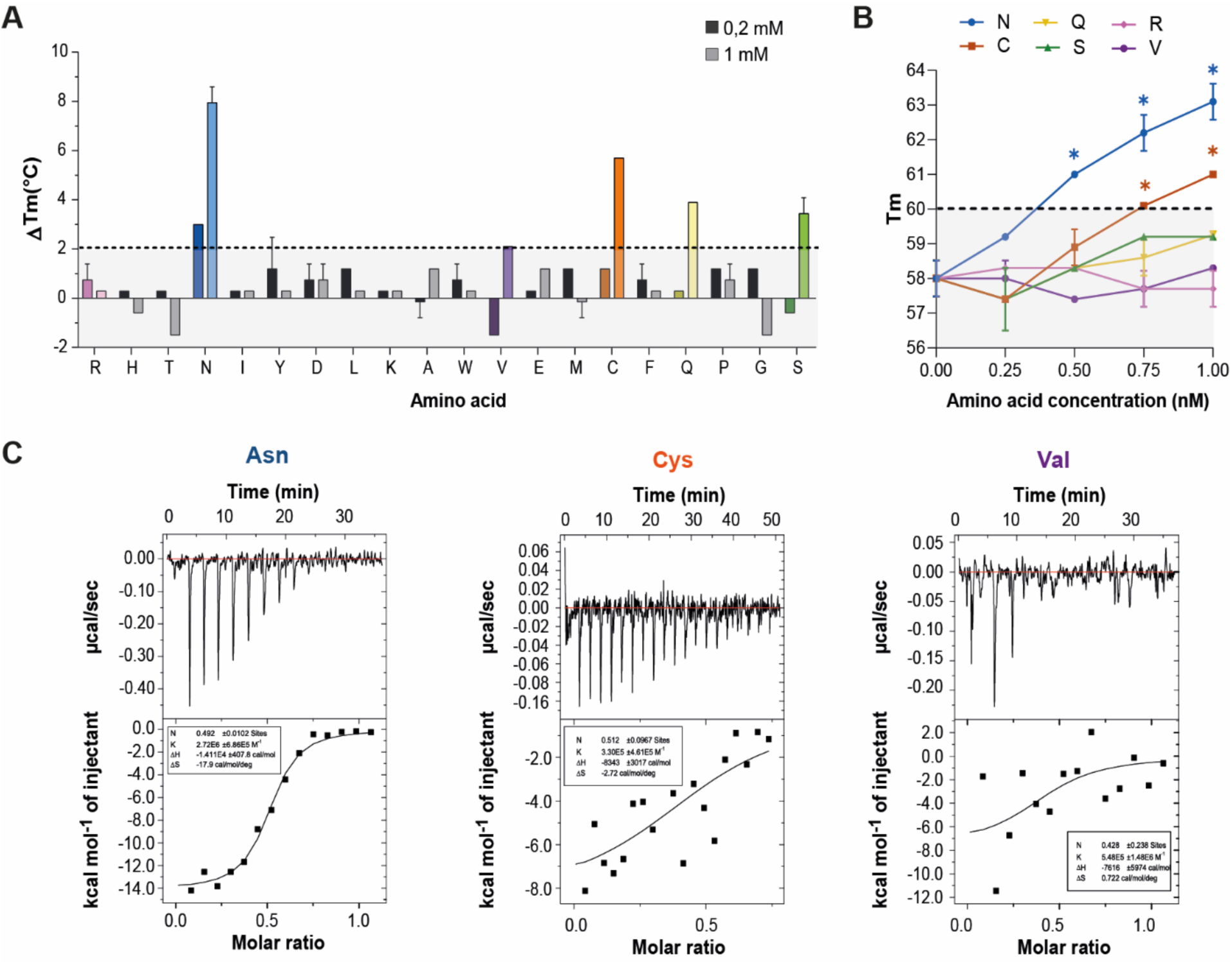
YhdW from *Citrobacter rodentium* binds asparagine. **(A)** Fluorescence thermal shift assay (FTSA) screening for ligands of *C. rodentium* YhdW. Purified YhdW-8His protein (1.5 µM) was incubated with a panel of amino acids at concentrations of 0.2 mM (dark color) or 1 mM (light color). Samples were subjected to a temperature gradient, and melting temperatures (Tm) were determined using SYPRO Orange fluorescence. Changes in Tm relative to the amino acid–free control are shown (ΔTm). **(B)** Dose-dependent FTSA analysis. Purified YhdW-8His (2.5 µM) was incubated with 0.25, 0.5, 0.75, or 1 mM asparagine (Asn), cysteine (Cys), serine (Ser), glutamine (Gln), arginine (Arg), or valine (Val). Tm values were determined based on SYPRO Orange fluorescence. Significant differences between conditions with and without amino acids are indicated by * (*p* < 0.05, ANOVA test, n = 3). The dotted line indicates the 2°C threshold for ligand binding. **(C)** Isothermal titration calorimetry (ITC) analysis of purified YhdW-8His. Asn, Val, or Cys were titrated into apo-YhdW (200 µM). Upper panel: raw data. Lower panel: integrated heat changes plotted as a function of the amino acid/YhdW molar ratio. The solid line represents the best fit to a one-site binding model. Each experiment was repeated four times; a representative binding isotherm is shown.

Isothermal titration calorimetry (ITC) further confirmed specific binding of asparagine, with a dissociation constant (Kd) of 0.37 µM. No binding was detected for valine, while cysteine showed weak and non-specific interactions (Figure 5C). These results demonstrate that YhdW from *C. rodentium* is a high-affinity asparagine-binding protein.

### YhdWXYZ is not a primary Asn importer in *C. rodentium*

To investigate the physiological role of YhdWXYZ, growth of the Δ*yhdWXYZ* mutant from *C. rodentium* was analyzed in minimal medium. No growth defect was observed compared to the WT strain (Figure 5A). Asparagine can be synthesized endogenously by two asparagine synthetases encoded by *asnA* and *asnB* genes. Asparagine auxotrophy was induced using a Δ*asnAΔasnB* mutant. Supplementation with 100 µM asparagine did not restore growth of the auxotrophic strains, whereas 1 mM asparagine restored growth without differences between the *ΔasnAΔasnB* and the *ΔasnAΔasnBΔyhdWXYZ* mutant strains (Figure 6A-6B).

**Figure 6.**
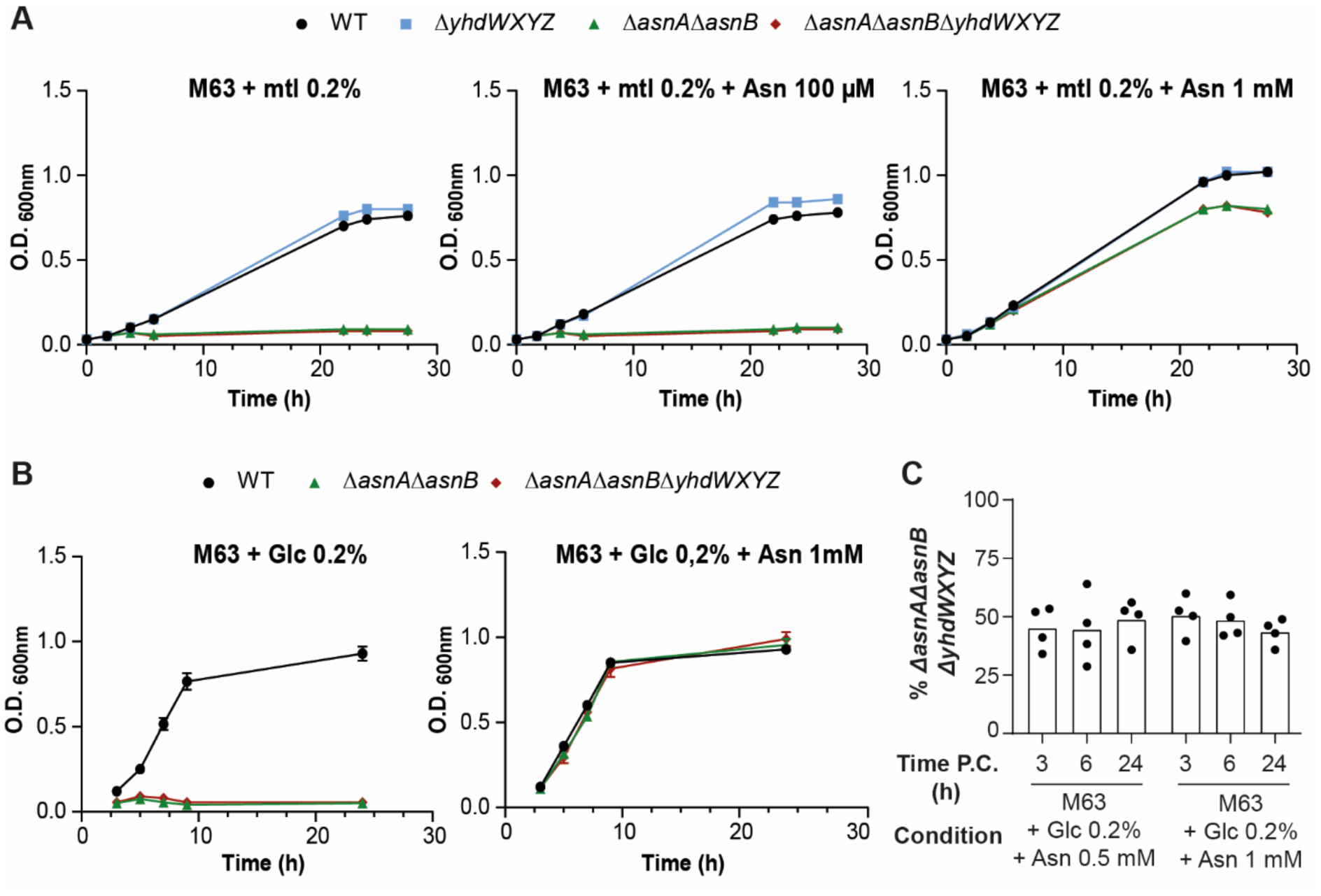
YhdWXYZ from *Citrobacter rodentium* is not a primary asparagine transporter. **(A)** Growth curves of the wild-type (WT) strain of *C. rodentium* and the isogenic mutants Δ*yhdWXYZ*, Δ*asnAΔasnB*, and Δ*asnAΔyhdWXYZΔasnB*. Strains were grown in M63 medium supplemented with 0.2% mannitol (left), or with 100 µM (center) or 1 mM (right) asparagine (Asn). Experiments were performed four times; a representative growth curve is shown. **(B)** Growth curves of the WT strain and the isogenic mutants Δ*asnAΔasnB* and Δ*asnAΔyhdWXYZΔasnB* grown in M63 medium supplemented with 0.2% glucose (left) or 1 mM Asn (right). Data are representative of two independent experiments (N = 2). **(C)** Competition assay. The Δ*asnAΔasnB* strain and the isogenic Δ*asnAΔasnBΔyhdWXYZ*-KanR mutant were co-cultured at a 1:1 ratio in M63 medium supplemented with 0.2% glucose and either 500 µM or 1 mM Asn. Colony-forming units (CFUs) were determined after 3, 6, and 24 h. Results are expressed as the percentage of the triple mutant relative to the total population: (CFU Δ*asnAΔasnBΔyhdWXYZ* / total CFU) × 100. Data represent the mean of four biological replicates (N = 4).

However, the absence of *yhdWXYZ* may lead to a marginally diminished growth rate. In order to address this point, co-cultures of the Δ*asnA*Δ*asnB* and Δ*asnA*Δ*asnB*Δ*yhdWXYZ* -Kan^R^ mutant strains were performed in a 1:1 proportion in M63 medium containing 500 µM or 1 mM Asn. CFU enumeration was conducted at 2-, 6- or 24 hours post-inoculation. At each time point, the number of CFU for both strains was comparable (Figure 5C). The two strains exhibited equivalent levels of fitness (Figure 6C) indicating that YhdWXYZ is not a primary Asn transporter in *C. rodentium* in these conditions of culture and that its function is redundant with yet unknown transporters.

### YhdWXYZ contributes to virulence *in vivo*

The pathogen *C. rodentium* shares key virulence mechanisms with EPEC and EHEC, particularly the LEE-encoded type III secretion system, which enables the bacterium to inject effector proteins into host epithelial cells and induce characteristic attaching-and-effacing (A/E) lesions. Because of these conserved mechanisms, *C. rodentium* is widely used as an *in vivo* model for studying EPEC- and EHEC-driven intestinal pathogenesis (Collins *et al*., 2014).

To assess the role of YhdWXYZ in host colonization, mice were infected with WT, Δ*yhdW*, or Δ*yhdWXYZ* strains of *C. rodentium*. Body weight remained similar across groups throughout the experiment (Figure 7A). However, bacterial loads in feces were significantly lower in mice infected with mutant strains at early time points (4–6 days post-infection), indicating reduced colonization efficiency (Figure 7B).

**Figure 7.**
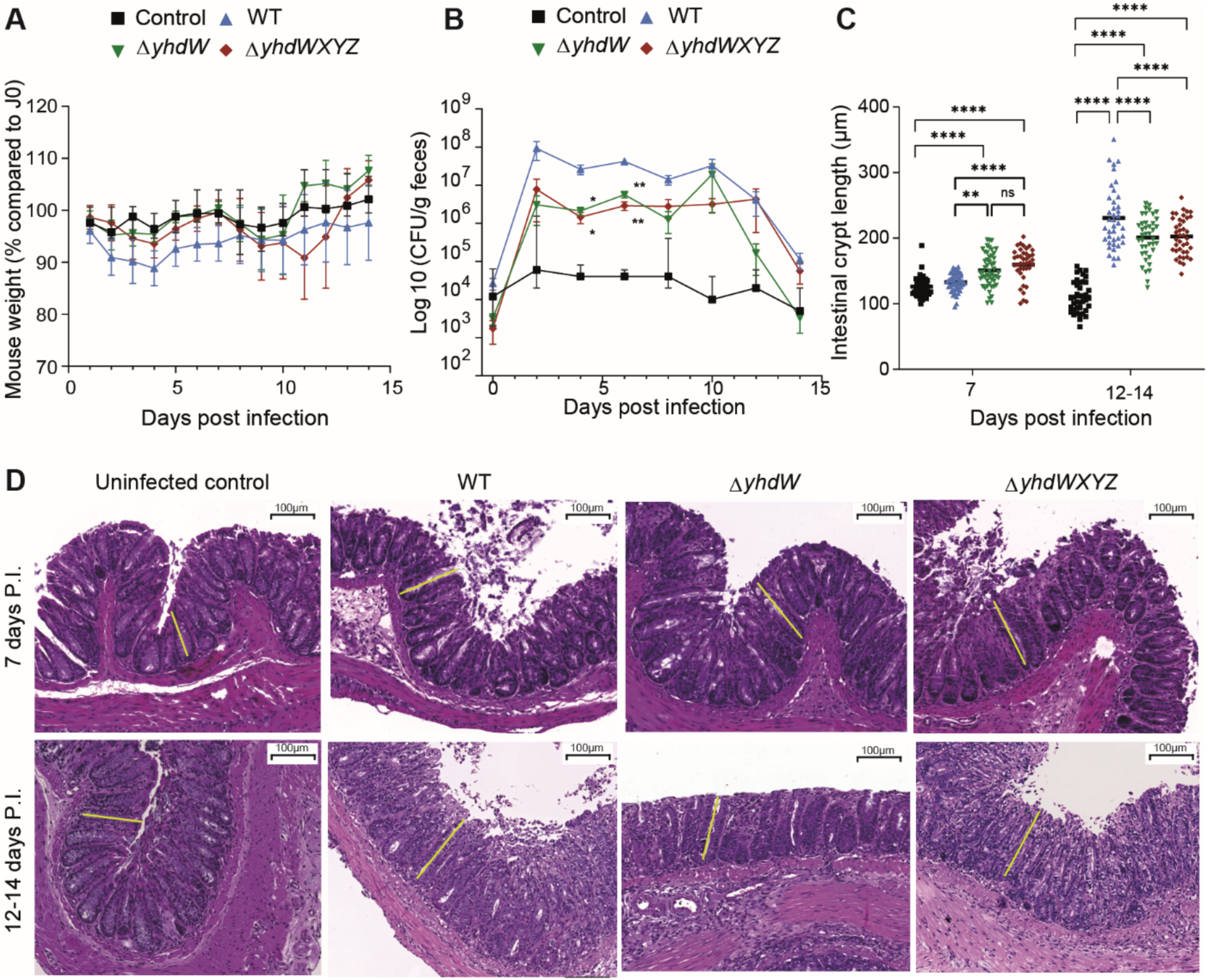
YhdWXYZ from *Citrobacter rodentium* contributes to virulence *in vivo*. C57BL/6 mice were orally infected with PBS (control) or with bacterial suspensions of *C. rodentium* WT, Δ*yhdW*, or Δ*yhdWXYZ* strains (n = 9 per group). **(A)** Mouse body weight was monitored daily. Data are expressed as the percentage of initial body weight at the time of infection. **(B)** Fecal samples were collected every two days, weighed, homogenized in PBS, and plated on MacConkey sorbitol agar. Bacterial loads were determined as colony-forming units (CFU) per gram of feces. Statistical analysis was performed using a mixed-effects model (REML) followed by Tukey’s multiple comparisons test (*P* < 0.05; \**P* < 0.01). **(C)** Mice were divided into two groups: one sacrificed at day 7 post-infection (n = 5) and the other between days 12 and 14 (n = 4). Colons were collected, fixed, paraffin-embedded, sectioned, and stained with hematoxylin and eosin (H&E). Crypt hyperplasia was quantified by measuring the length of 10 well-oriented colonic crypts per mouse. Statistical analysis was performed using a mixed-effects model (REML) followed by Tukey’s multiple comparisons test (ns, not significant; \**P* < 0.01; \*\*\**P* < 0.0001). **(D)** Representative H&E-stained distal colon sections from control, WT-, Δ*yhdW*-, and Δ*yhdWXYZ*-infected mice at 7 days post-infection (p.i.) and 12–14 days p.i. Scale bar, 100 µm. Crypt hyperplasia is indicated by a yellow bar.

Histological analysis of colon tissues revealed differences in crypt hyperplasia. At 7 days post-infection, mutants induced slightly increased hyperplasia compared to WT. In contrast, at later time points (12–14 days), hyperplasia was significantly reduced in mutant-infected mice (Figures 7C-7D). These results suggest that YhdWXYZ contributes to early colonization and persistence during infection.

## DISCUSSION

Based on sequence analysis, all components of the YhdWXYZ ABC transporter from *C. rodentium* were classified within the amino acid transporter family. Structural comparisons identified YhdW as most closely related to BztA from *Brucella ovis*, whose ligand specificity has been characterized in homologous systems from *Rhodobacter capsulatus* (Zheng & Haselkorn, 1996). The BztABCD transporter has been shown to support growth in ammonium-depleted conditions supplemented with specific amino acids, including glutamate, glutamine, aspartate, and asparagine.

Biochemical analyses demonstrated that YhdW binds asparagine with high affinity and specificity. Despite this, no growth defect was observed for the Δ*yhdWXYZ* mutant in minimal medium, even in strains unable to synthesize asparagine endogenously. These results strongly suggest the presence of alternative asparagine transport systems in *C. rodentium*.

YhdW and BztA are homologs of the AapJ SBP. In the symbiotic bacteria *Rhizobium leguminosarum,* AapJ was shown to be part of a L-amino acid transporter with broad specificity (Walshaw & Poole, 1996) and to be positively regulated by nitrogen limitation (Walshaw *et al*, 1997). In this organism, the Aap system plays a key role in nitrogen metabolism and symbiosis and contributes to both amino acid uptake and efflux. Mutants lacking Aap exhibit reduced persistence in host tissues despite retaining nitrogen fixation capacity (Lodwig *et al*., 2003). Similarly, homologous systems in *Brucella abortus* have been identified as virulence factors required for early stages of infection (Tian *et al*, 2018).

Consistent with these observations, our *in vivo* experiments demonstrate that YhdWXYZ contributes to host colonization in *C. rodentium*. Mutant strains exhibited reduced bacterial loads during early stages of infection, indicating impaired colonization efficiency. Interestingly, although bacterial clearance occurred with similar kinetics at later time points, histological analyses revealed altered patterns of crypt hyperplasia. These findings suggest that YhdWXYZ is not strictly required for virulence but plays a role in optimizing bacterial persistence and host interaction dynamics.

One possible explanation for this phenotype is a reduced capacity to acquire specific nutrients within the host environment. Although no clear role for YhdWXYZ in amino acid uptake was identified under laboratory conditions, nutrient availability *in vivo* is highly dynamic and shaped by host diet and microbiota composition (Krautkramer *et al*, 2021). Competition for amino acids is a key determinant of microbial fitness in the gut, and transport systems may provide a selective advantage during early colonization stages (Caballero-Flores *et al*, 2023).

To strengthen this hypothesis, a global study of the roles of ABC transporters was conducted in uropathogenic *E. coli* (UPEC) causing urinary tract infections (Shea *et al*, 2024). The growths of several mutants were tested in synthetic media as well as in human urine. The Δ*yhdWXYZ* mutant only presented a slightly reduced growth in urine. More interesting, this mutant competitive index was significantly lower than the wild type strain in the kidneys during infection. These results are consistent with ours in highlighting the importance of YhdWXYZ during host interaction and not in *in vitro* conditions.

Recent studies have highlighted the importance of amino acid metabolism in *C. rodentium* colonization, particularly under conditions of microbiota competition (Caballero-Flores *et al*, 2020) (Caballero-Flores *et al*, 2021). In this context, YhdWXYZ may contribute to nutrient acquisition under specific *in vivo* conditions that are not reproduced *in vitro*. Further work will be required to identify the physiological substrates and environmental conditions under which this transporter is active.

A key finding of this study is the functional divergence of YhdWXYZ between *E. coli* K-12 and *C. rodentium*. In *E. coli* K-12, the *yhdW* gene is a pseudogene, resulting in the absence of a functional SBP. Under these conditions, deletion of *yhdWXYZ* leads to increased intracellular accumulation of lomefloxacin and increased sensitivity to this antibiotic, suggesting a role in antibiotic efflux or homeostasis. This result is in line with previous work where the fitness of the complete single gene knock-out mutant library of *E. coli* K-12 was evaluated in presence of 114 stressors (antibiotics, detergents, drugs…). Ciprofloxacin, Norfloxacin (fluoroquinolones) and mecillinam were found to affect the fitness of the *yhdZ* mutant (Nichols *et al*., 2011).

This observation raises the possibility that, in the absence of the SBP, the transporter operates with altered substrate specificity or directionality. Although ABC transporters are generally considered unidirectional, several studies have demonstrated bidirectional transport capabilities. For example, the Aap system from *R. leguminosarum* has been shown to mediate both import and export of amino acids (Hosie *et al*, 2001).

Our findings suggest that YhdWXYZ may adopt distinct functional roles depending on its structural composition. In *C. rodentium*, the complete transporter primarily supports host adaptation, whereas in *E. coli* K-12, the absence of the SBP is associated with a shift toward antibiotic homeostasis. This functional plasticity highlights the evolutionary flexibility of ABC transporters and underscores their capacity to adapt to different ecological niches and selective pressures.

Overall, this study reveals that the presence or absence of a single component, the SBP, can profoundly alter the physiological role of an ABC transporter. These findings provide new insights into the adaptability of bacterial transport systems and their contribution to both antibiotic resistance and host–pathogen interactions.

## ACKNOWLEDMENTS

We would like to pay tribute to the memory of our colleague, Michel Droux, whose expertise in biochemistry made a significant contribution to this study. We thank Olivier Espéli for critical reading of the manuscript. The authors would like to thank the staff of the Animalerie Dubois, Julie Ulmann, Angeline Clair et Laetitia Averty, for their excellent animal care and technical support.

CB was supported by a grant from the French Ministry of Higher Education and Research. CR and AR received a grant from FR Bioenvis. Finally, we acknowledge the support of the BF2i and MAP laboratories for the use of their experimental facilities.

## SUPPLEMENTARY FIGURES

**Figure S1:**
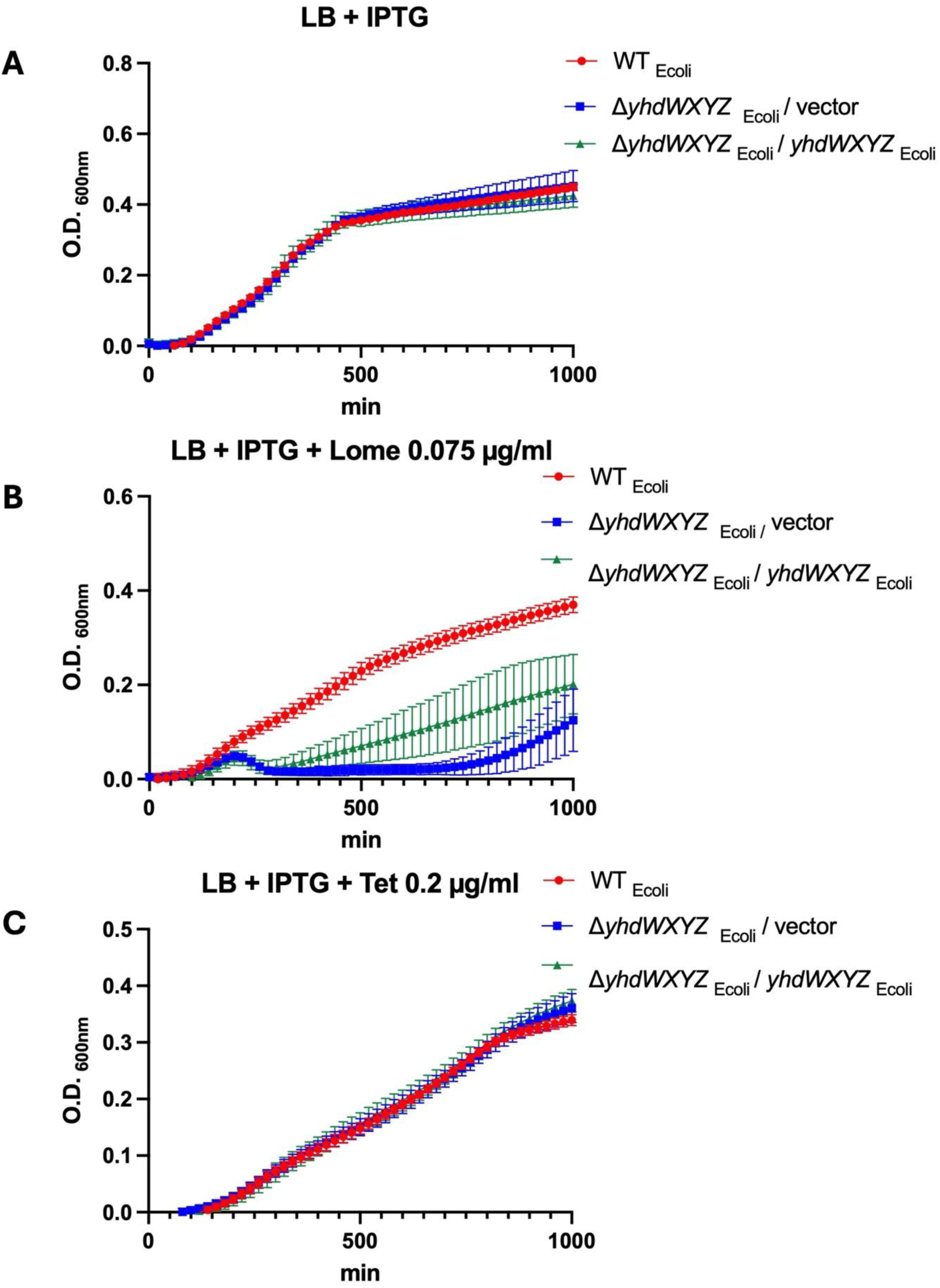
Growth of *E.coli* K-12 challenged with antibiotics. Growth curves of the WT strain (A), the *L’lyhdWXYZ* mutant / vector (B) and the *L’lyhdWXYZ I yhdWXYZ* strain (C). The bacteria were grown), at 37°C in LB+ IPTG lmM, or in LB + IPTG lmM + Lomefloxacin (0.075 µg/rnL) or Tetracycline (0.2 µg / rnL). The O.D._60onm_ was measured every 20 min. The mean and SD of 7 replicates are shown.

**Figure S2:**
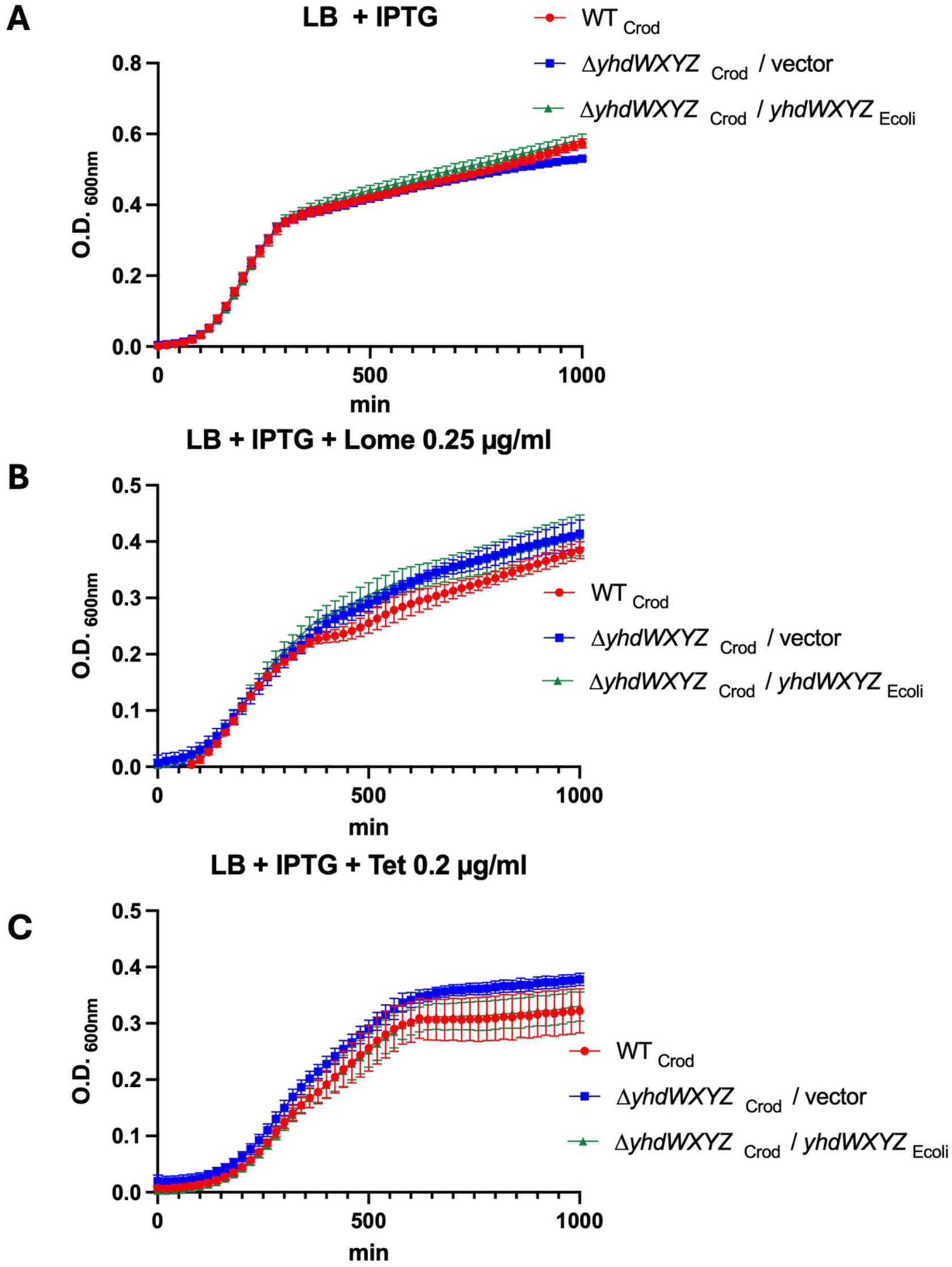
Growth of *C. rodentium* challenged with antibiotics. Growth curves of *C. rodentium* RCL2 WT strain (A), the *tiyhdWXYZ isogenic* mutant *I* vector (B) and the ti *yhdWXYZ mutant I* complemented by *yhdWXYZ* of *E. coli* K-12 (C). The bacteria were grown in LB + IPTG lmM, or in LB +lmM IPTG + subinhibitory concentrations of: Lomefloxacin (0.25 µg/mL) or Tetracycline (0.2 µg / mL). The O.D._600nm_ was measured every 20 min at 37°C. The mean and SD of 7 replicates are shown.

**Figure S3.**
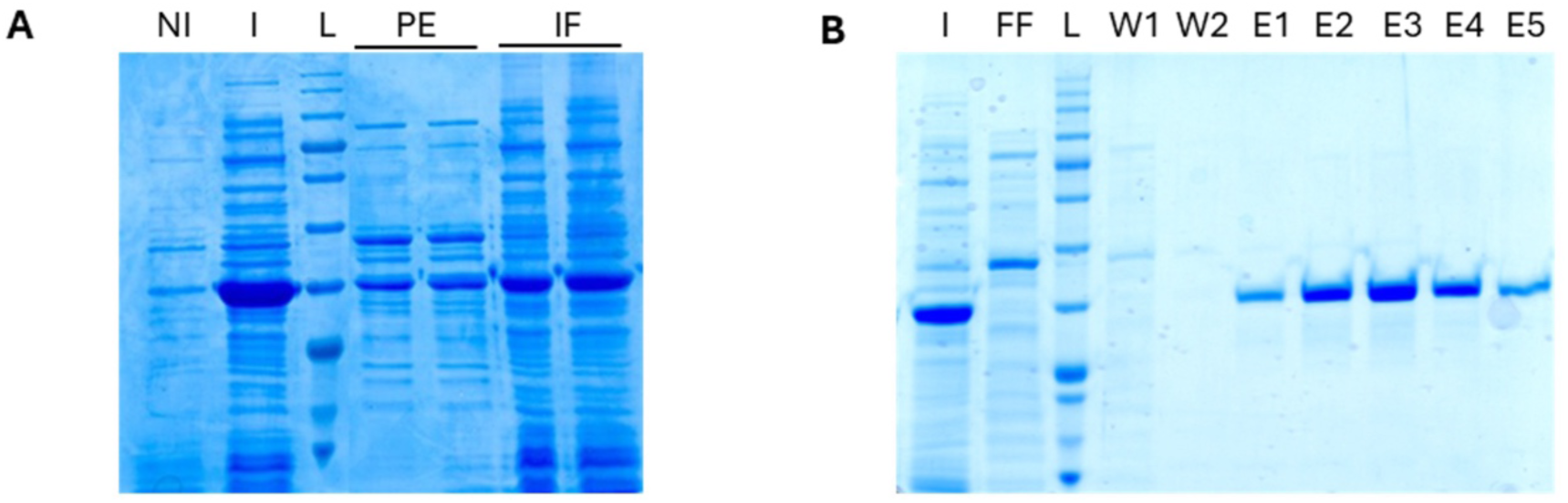
: Production and purification of C. rodentium YhdW. (A) Surproduction of His-tagged YhdW. NI: non induced crude extract; I: induced crude extract; L: ladder; PE: periplasmic extract; IF: insoluble fraction. (B) Ni-NTA Purification. I: induced crude extract; FF: Flow through; L: ladder; W1: 10 mM imidazole wash; W2: 30 mM imidazole wash, E1-E5: 150 mM imidazole elution.

**Table S1:** Strains used in this study.

| Strains | Characteristics | Source and references |
| --- | --- | --- |
| W3110 | *Escherichia coli* K12 (F- mcrAmcrB IN (rrnD-rrnE)1 λ-) | Lab collection |
| W3110 $\Delta yhdWXYZ$ | W3110, $\Delta yhdWXYZ$ | This work |
| RCL2 | NalR strain derived from *Citrobacter rodentium* DBS100 (ATCC 51459) | (Gueguen & Cascales, 2013) |
| RCL2 $\Delta yhdWXYZ$ | RCL2, $\Delta yhdWXYZ$ | This work |
| RCL2 $\Delta asnAasnB$ | RCL2, $\Delta asnA$, $\Delta asnB$ | This work |
| RCL2 $\Delta asnAasnB yhdWXYZ$ | RCL2, $\Delta asnA$, $\Delta asnB$, $\Delta yhdWXYZ$ | This work |
| BL21 | *Escherichia coli* B; F- ompT gal [dcm] [lon] hsdSB (rB- mB-) (DE3) | (Studier & Moffatt, 1986) |

**Table S2:** Plasmids used in this study.

| Plasmids | Characteristics | Source and references |
| --- | --- | --- |
| pKOBEG | CmR, arabinose-inducible λred $\gamma\beta\alpha$ operon, oriR101, repA101-ts | (Chaveroche *et al*, 2000) |
| pKD46 | AmpR, arabinose-inducible λred $\gamma\beta\alpha$ operon, oriR101, repA101-ts | (Datsenko & Wanner, 2000) |
| pKD3 | AmpR, CmR gene flanked by FRT sites, oriRγR6k | (Datsenko & Wanner, 2000) |
| pLux | KanR, *luxCDABE*, Ori p15A | (Cayron *et al*, 2017) |
| P$yhdW::lux$ | The *E.coli* W3110 *yhdW* promoter cloned upstream of the *luxCDABE* genes in pLux | This work |
| P$yhdX::lux$ | The *E.coli* W3110 *yhdX* promoter cloned upstream of the *luxCDABE* genes in pLux | This work |
| pET30 | KanR, T7 promoter, overexpression vector | Lab collection |
| pET30-YhdW-HT | *C.rodentium* RCL2 *yhdW* gene fused to C-ter 8 His-Tag, cloned in pET30 | This work |
| pVLT33 | KanR, tac promoter, RSF1010-ORIV | Lab collection |
| pVLT33<br>yhdWXYZ | The yhdW promoter, yhdW pseudogene, yhdX promoter and yhdXYZ genes of *E.coli* cloned in pVLT33 | This work |

**Table S3:**
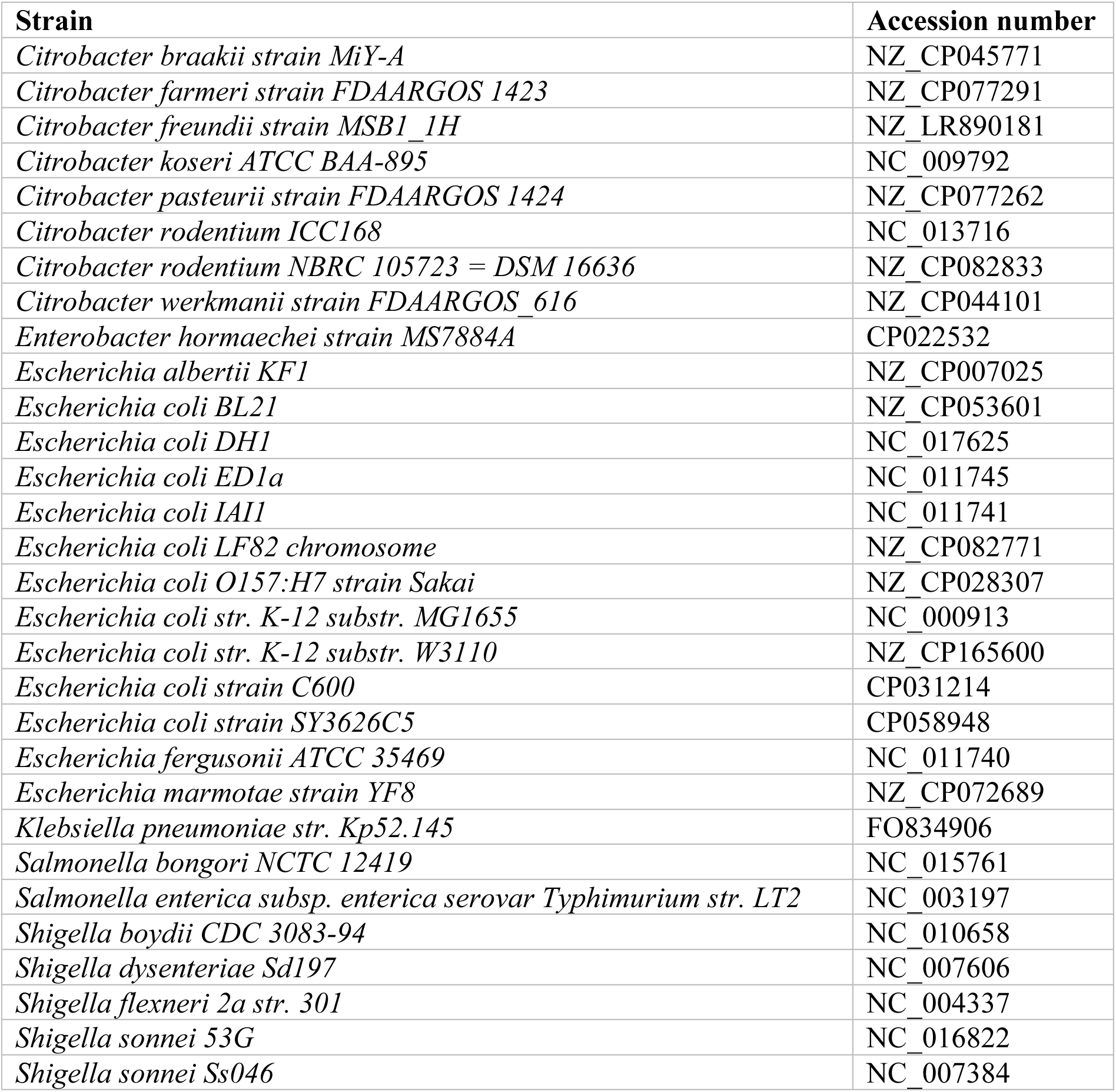
List of genome accession numbers.

| Strain | Accession number |
| --- | --- |
| *Citrobacter braakii* strain MiY-A | NZ_CP045771 |
| *Citrobacter farmeri* strain FDAARGOS 1423 | NZ_CP077291 |
| *Citrobacter freundii* strain MSB1_1H | NZ_LR890181 |
| *Citrobacter koseri* ATCC BAA-895 | NC_009792 |
| *Citrobacter pasteurii* strain FDAARGOS 1424 | NZ_CP077262 |
| *Citrobacter rodentium* ICC168 | NC_013716 |
| *Citrobacter rodentium* NBRC 105723 = DSM 16636 | NZ_CP082833 |
| *Citrobacter werkmanii* strain FDAARGOS_616 | NZ_CP044101 |
| *Enterobacter hormaechei* strain MS7884A | CP022532 |
| *Escherichia albertii* KF1 | NZ_CP007025 |
| *Escherichia coli* BL21 | NZ_CP053601 |
| *Escherichia coli* DH1 | NC_017625 |
| *Escherichia coli* ED1a | NC_011745 |
| *Escherichia coli* IAI1 | NC_011741 |
| *Escherichia coli* LF82 chromosome | NZ_CP082771 |
| *Escherichia coli* O157:H7 strain Sakai | NZ_CP028307 |
| *Escherichia coli* str. K-12 substr. MG1655 | NC_000913 |
| *Escherichia coli* str. K-12 substr. W3110 | NZ_CP165600 |
| *Escherichia coli* strain C600 | CP031214 |
| *Escherichia coli* strain SY3626C5 | CP058948 |
| *Escherichia fergusonii* ATCC 35469 | NC_011740 |
| *Escherichia marmotae* strain YF8 | NZ_CP072689 |
| *Klebsiella pneumoniae* str. Kp52.145 | FO834906 |
| *Salmonella bongori* NCTC 12419 | NC_015761 |
| *Salmonella enterica* subsp. *enterica* serovar *Typhimurium* str. LT2 | NC_003197 |
| *Shigella boydii* CDC 3083-94 | NC_010658 |
| *Shigella dysenteriae* Sd197 | NC_007606 |
| *Shigella flexneri* 2a str. 301 | NC_004337 |
| *Shigella sonnei* 53G | NC_016822 |
| *Shigella sonnei* Ss046 | NC_007384 |

## REFERENCES

Abramson J, Adler J, Dunger J, Evans R, Green T, Pritzel A, Ronneberger O, Willmore L, Ballard AJ, Bambrick J et al (2024) Accurate structure prediction of biomolecular interactions with AlphaFold 3. Nature 630: 493–500

Caballero-Flores G, Pickard JM, Fukuda S, Inohara N, Nunez G (2020) An Enteric Pathogen Subverts Colonization Resistance by Evading Competition for Amino Acids in the Gut. Cell Host Microbe 28: 526–533 e525

Caballero-Flores G, Pickard JM, Nunez G (2021) Regulation of *Citrobacter rodentium* colonization: virulence, immune response and microbiota interactions. Curr Opin Microbiol 63: 142–149

Caballero-Flores G, Pickard JM, Núñez G (2023) Microbiota-mediated colonization resistance: mechanisms and regulation. Nature Reviews Microbiology 21: 347–360

Cayron J, Prudent E, Escoffier C, Gueguen E, Mandrand-Berthelot MA, Pignol D, Garcia D, Rodrigue A (2017) Pushing the limits of nickel detection to nanomolar range using a set of engineered bioluminescent *Escherichia coli*. Environ Sci Pollut Res Int 24: 4–14

Collins JW, Keeney KM, Crepin VF, Rathinam VA, Fitzgerald KA, Finlay BB, Frankel G (2014) Citrobacter rodentium: infection, inflammation and the microbiota. Nat Rev Microbiol 12: 612–623

Crepin VF, Habibzay M, Glegola-Madejska I, Guenot M, Collins JW, Frankel G (2015) Tir Triggers Expression of CXCL1 in Enterocytes and Neutrophil Recruitment during *Citrobacter rodentium* Infection. Infect Immun 83: 3342–3354

Datsenko KA, Wanner BL (2000) One-step inactivation of chromosomal genes in *Escherichia coli* K-12 using PCR products. Proc Natl Acad Sci U S A 97: 6640–6645

Davidson AL, Dassa E, Orelle C, Chen J (2008) Structure, function, and evolution of bacterial ATP-binding cassette systems. Microbiol Mol Biol Rev, pp. 317–364

Davies, J. S., Currie, M. J., Wright, J. D., Newton-Vesty, M. C., North, R. A., Mace, P. D., et al. (2021) Selective Nutrient Transport in Bacteria: Multicomponent Transporter Systems Reign Supreme In Front Mol Biosci, 8: 699222.

de Boer M, Gouridis G, Vietrov R, Begg SL, Schuurman-Wolters GK, Husada F, Eleftheriadis N, Poolman B, McDevitt CA, Cordes T (2019) Conformational and dynamic plasticity in substrate-binding proteins underlies selective transport in ABC importers. In: Elife 8:e44652

Holm L, Laiho A, Toronen P, Salgado M (2023) DALI shines a light on remote homologs: One hundred discoveries. Protein Sci 32: e4519

Hosie AH, Allaway D, Jones MA, Walshaw DL, Johnston AW, Poole PS (2001) Solute-binding protein-dependent ABC transporters are responsible for solute efflux in addition to solute uptake. Mol Microbiol 40: 1449–1459

Jago MJ, Soley JK, Denisov S, Walsh CJ, Gifford DR, Howden BP, Lagator M (2025) High-throughput method characterizes hundreds of previously unknown antibiotic resistance mutations. Nat Commun 16: 780

Krautkramer KA, Fan J, Backhed F (2021) Gut microbial metabolites as multi-kingdom intermediates. Nat Rev Microbiol 19: 77–94

Lodwig EM, Hosie AH, Bourdes A, Findlay K, Allaway D, Karunakaran R, Downie JA, Poole PS (2003) Amino-acid cycling drives nitrogen fixation in the legume-Rhizobium symbiosis. Nature 422: 722–726

Moussatova A, Kandt C, O’Mara ML, Tieleman DP (2008) ATP-binding cassette transporters in *Escherichia coli*. In: Biochim Biophys Acta, 1778: 1757–1771.

Nichols RJ, Sen S, Choo YJ, Beltrao P, Zietek M, Chaba R, Lee S, Kazmierczak KM, Lee KJ, Wong A et al (2011) Phenotypic landscape of a bacterial cell. Cell 144: 143–156

Rome K, Borde C, Taher R, Cayron J, Lesterlin C, Gueguen E, De Rosny E, Rodrigue A (2018) The Two-Component System ZraPSR Is a Novel ESR that Contributes to Intrinsic Antibiotic Tolerance in *Escherichia coli*. J Mol Biol 430: 4971–4985

Shea AE, Forsyth VS, Stocki JA, Mitchell TJ, Frick-Cheng AE, Smith SN, Hardy SL, Mobley HLT (2024) Emerging roles for ABC transporters as virulence factors in uropathogenic *Escherichia coli*. Proc Natl Acad Sci U S A 121: e2310693121

Sun R, Zhao X, Meng Q, Huang P, Zhao Q, Liu X, Zhang W, Zhang F, Fu Y (2022) Genome-Wide Screening and Characterization of Genes Involved in Response to High Dose of Ciprofloxacin in *Escherichia coli*. Microb Drug Resist 28: 501–510

Thomas C, Aller SG, Beis K, Carpenter EP, Chang G, Chen L, Dassa E, Dean M, Duong Van Hoa F, Ekiert D et al (2020) Structural and functional diversity calls for a new classification of ABC transporters. FEBS Lett 594: 3767–3775

Thomas C, Tampe R (2020) Structural and Mechanistic Principles of ABC Transporters. Annu Rev Biochem 89: 605–636

Tian M, Bao Y, Li P, Hu H, Ding C, Wang S, Li T, Qi J, Wang X, Yu S (2018) The putative amino acid ABC transporter substrate-binding protein AapJ2 is necessary for *Brucella* virulence at the early stage of infection in a mouse model. Vet Res 49: 32

Vergalli J, Dumont E, Pajovic J, Cinquin B, Maigre L, Masi M, Refregiers M, Pages JM (2018) Spectrofluorimetric quantification of antibiotic drug concentration in bacterial cells for the characterization of translocation across bacterial membranes. Nat Protoc 13: 1348–1361

Walshaw DL, Poole PS (1996) The general L-amino acid permease of *Rhizobium leguminosarum* is an ABC uptake system that also influences efflux of solutes. Mol Microbiol 21: 1239–1252

Walshaw DL, Reid CJ, Poole PS (1997) The general amino acid permease of *Rhizobium leguminosarum* strain 3841 is negatively regulated by the Ntr system. FEMS Microbiol Lett 152: 57–64

Yu F, Li X, Wang F, Liu Y, Zhai C, Li W, Ma L, Chen W (2023) TLTC, a T5 exonuclease-mediated low-temperature DNA cloning method. Front Bioeng Biotechnol 11: 1167534

Zheng S, Haselkorn R (1996) A glutamate/glutamine/aspartate/asparagine transport operon in *Rhodobacter capsulatus*. Mol Microbiol 20: 1001–1011

## References

Cayron J, Prudent E, Escoffier C, Gueguen E, Mandrand-Berthelot MA, Pignol D, Garcia D, Rodrigue A (2017) Pushing the limits of nickel detection to nanomolar range using a set of engineered bioluminescent Escherichia coli. Environ Sci Pollut Res Int 24: 4–14

Chaveroche MK, Ghigo JM, d’Enfert C (2000) A rapid method for efficient gene replacement in the filamentous fungus Aspergillus nidulans. In: Nucleic Acids Res, p. E97.

Datsenko KA, Wanner BL (2000) One-step inactivation of chromosomal genes in Escherichia coli K-12 using PCR products. In: Proc Natl Acad Sci U S A, pp. 6640–6645.

Gueguen E, Cascales E (2013) Promoter swapping unveils the role of the Citrobacter rodentium CTS1 type VI secretion system in interbacterial competition. In: Appl Environ Microbiol, pp. 32–38.

Studier FW, Moffatt BA (1986) Use of bacteriophage T7 RNA polymerase to direct selective high-level expression of cloned genes. In: J Mol Biol, pp. 113–130.

